# Multiple requirements for Rab GTPases in the development of *Drosophila* tracheal dorsal branches and terminal cells

**DOI:** 10.1101/873232

**Authors:** Benedikt T. Best, Maria Leptin

## Abstract

The tracheal epithelium in fruit fly larvae is a popular model for multi- and unicellular migration and morphogenesis. Like all epithelial cells, tracheal cells use Rab GTPases to organise their internal membrane transport, resulting in the specific localisation or secretion of proteins on the apical or basal membrane compartments. Some contributions of Rabs to junctional remodelling and governance of tracheal lumen contents are known, but it is reasonable to assume that they play important further roles in morphogenesis. This pertains in particular to terminal tracheal cells, specialised branch-forming cells that drastically reshape both their apical and basal membrane during the larval stages. We performed a loss-of-function screen in the tracheal system, knocking down endogenously tagged alleles of 26 Rabs by targeting the tag via RNAi. This revealed that at least 14 Rabs are required to ensure proper cell fate specification and migration of the dorsal branches, as well as their epithelial fusion with the contralateral dorsal branch. The screen implicated four Rabs in the subcellular morphogenesis of terminal cells themselves. Further tests suggested residual gene function after knockdown, leading us to discuss the limitations of this approach. We conclude that more Rabs than identified here may be important for tracheal morphogenesis, and that the tracheal system offers great opportunities for studying several Rabs that have barely been characterised so far.

## Introduction

The larvae of *Drosophila melanogaster* breathe through a network of tracheal tubes reminiscent of vertebrate blood vessels. The anatomy of the tracheal system is defined during the second half of embryogenesis, and originates from ten bilateral ectoderm-derived tracheal placodes. Tracheal cells migrate outwards from each placode in response to the fibroblast growth factor (FGF) Branchless (Bnl), secreted by small groups of cells around each placode (Du et al., 2017; Ohshiro et al., 2002; Sutherland et al., 1996). Most migratory movements of tracheal cells throughout embryogenesis are thus directed by the FGF receptor Breathless (Btl). During their movements, tracheal cells encounter signalling factors secreted by other tissues. In response to these positional cues, the cells express different transcription factors that in turn modify their morphogenetic behaviour. This leads to the adoption of genetic fates corresponding to the anatomical location of each cell within the tracheal system (Rao et al., 2015). Fusion events between primary branches from neighbouring tracheal placodes establish an interconnected tubular system by the end of embryogenesis, which then remains unchanged in its anatomy throughout the larval stages.

The dorsal branches have received much attention because their migratory movement and the mechanisms that determine cell fates within them are similar to vertebrate sprouting angiogenesis (Kotini et al., 2018). Dorsal branches supply the muscles of the dorsal body wall with oxygen and connect directly to the dorsal trunks, the largest tracheal tubes. Each body segment contains one pair of bilateral dorsal branches that meet at the dorsal midline near the end of embryogenesis. Each dorsal branch has one specialised fusion cell at the tip, which establishes contact with its contralateral counterpart and mediates the epithelial fusion of the two dorsal branches. Distal to this anastomosis, each dorsal branch has one terminal cell that attaches to a muscle, which it will later supply with oxygen. After the fully developed larva hatches, the anatomy of the tracheal system remains unchanged. Instead, the terminal cells located at branch tips throughout the network undergo subcellular branching morphogenesis, forming long cellular processes that wrap around oxygen-demanding tissues. Terminal cell branches contain subcellular tubes that are continuous with the tracheal lumen and deliver air to the target tissue.

Most tracheal tubes consist of a single layer of squamous epithelial cells, lined by a basal lamina towards the interior of the animal. On their lumenal membrane, tracheal cells secrete an apical extracellular matrix, which confers mechanical stability to the tubes and enables them to conduct air to the body’s tissues. This matrix strongly autofluoresces when illuminated with near-ultraviolet light (Lin et al., 2008), which we exploit here to image tracheal tubes in *Drosophila* larvae. Like all epithelial cells, tracheal cells use Rab GTPases to organise the delivery of proteins and membrane to specific membrane compartments. Rab proteins recruit various effectors, including key components of the vesicle trafficking machinery, such as kinesins and myosins (Campa and Hirsch, 2017), as well as tethering complexes (Cai et al., 2007). With their role as key regulators of membrane trafficking Rab GTPases are thus a promising group of genes to study for understanding tracheal morphogenesis. Some Rabs have been connected with specific roles in this developmental process. Rab11 is important for recycling E-cadherin and for junctional remodelling during branch migration and during cell intercalation, when branches transition from multicellular to unicellular architectures (Cheshire et al., 2008; Kerman et al., 2008). Rab9 and Rab5 are involved in tracheal development but not morphogenesis *per se*. Some functions of Rab5, Rab7 and Rab39, Rab10 and Rab11, as well as Rab35 have been identified in the specialised cells found at the tips of the dorsal branches, terminal cells and fusion cells, both of which undergo striking shape changes in the late stages of embryonic development (Caviglia et al., 2016; Dong et al., 2013; Jones et al., 2014; Schottenfeld-Roames et al., 2014; Schottenfeld-Roames and Ghabrial, 2012; Tsarouhas et al., 2007).

Tracheal cells’ response to signalling cues from other tissues and the conserved function of Rabs suggest that the receptors of these signals require Rabs for their delivery to the plasma membrane, as well as their recycling, degradation, and possibly as part of their activation mechanism. Since all signals come from surrounding tissues, the membrane surface where receptors must be located to sense them is the basal membrane domain. Alternatively, signalling factors could be endocytosed and meet their receptor in endosomal membrane compartments. In case of terminal cells and fusion cells, the known involvements of some Rabs in their subcellular morphogenesis suggested that these and other Rabs might play so far unknown roles in cell shape determination. We therefore conducted an RNAi-mediated knockdown screen of 27 *Drosophila rab* genes and looked for phenotypes relating to the dorsal branches and terminal cells in wandering third-instar larvae. This represents the endpoint of tracheal development before metamorphosis, during which most of the architecture is replaced by new tracheal cells (Djabrayan et al., 2014). To increase confidence in our results and eliminate false positives, we did not use RNAi transgenes directly targeting the Rabs but instead used a collection of endogenously YFP-tagged Rabs (Dunst et al., 2015) and knocked them down using an RNAi transgene targeting the tag. This approach is similar to the previously reported “*in vivo* GFP RNAi” (iGFPi) and “tag-mediated loss-of-function” methods (Neumüller et al., 2012; Pastor-Pareja and Xu, 2011). To reflect the tag and target involved, we term the method “YRab-YFPi” in this paper.

We explored knock down approaches targeting the tag at RNA and protein level, and tested their ability to identify maternally contributed gene products. We discuss advantages and limitations of this screening approach. Our screen results show that 14 Rabs are required in dorsal branch morphogenesis, with knockdown phenotypes indicating that these Rabs are involved in cell fate specification and/or epithelial fusion. Furthermore, Rab2, Rab6, Rab8 and Rab10 are required for the morphogenesis of terminal cells during the larval stages.

## Results

### Expression of Rabs in terminal tracheal cells

As a basis for screening the functions of Rabs, we first characterised their expression and compared the efficiency and validity of different ways of reducing gene function in terminal cells. To characterise Rab expression in terminal cells, we imaged L3 wandering larvae carrying endogenously YFP-tagged *rab* alleles (YRabs) and examined either YFP fluorescence in heat-fixed larvae, or dissected larvae and quantified the intensity of staining using an anti-GFP antibody (Fig. 1A-C). We distinguish three classes: YRabs that can be clearly detected by the YFP fluorescence in live or heat-fixed larvae (YRab1, 2, 6, 7, 11; Fig. 1D-G show stained specimens), YRabs that can be detected only by immunostaining (YRab5, 8, 10, 39, 40) and YRabs whose expression, if it exists, is too low to be distinguished from unspecific staining by either method (YRab3, 4, 9, 14, 18, 19, 21, 23, 26, 27, 30, 32, 35, X1, X4, X5, X6). All detectable YRabs showed a punctate staining in the TC cytoplasm with no particular enrichment anywhere in the cell.

**Figure 1.**
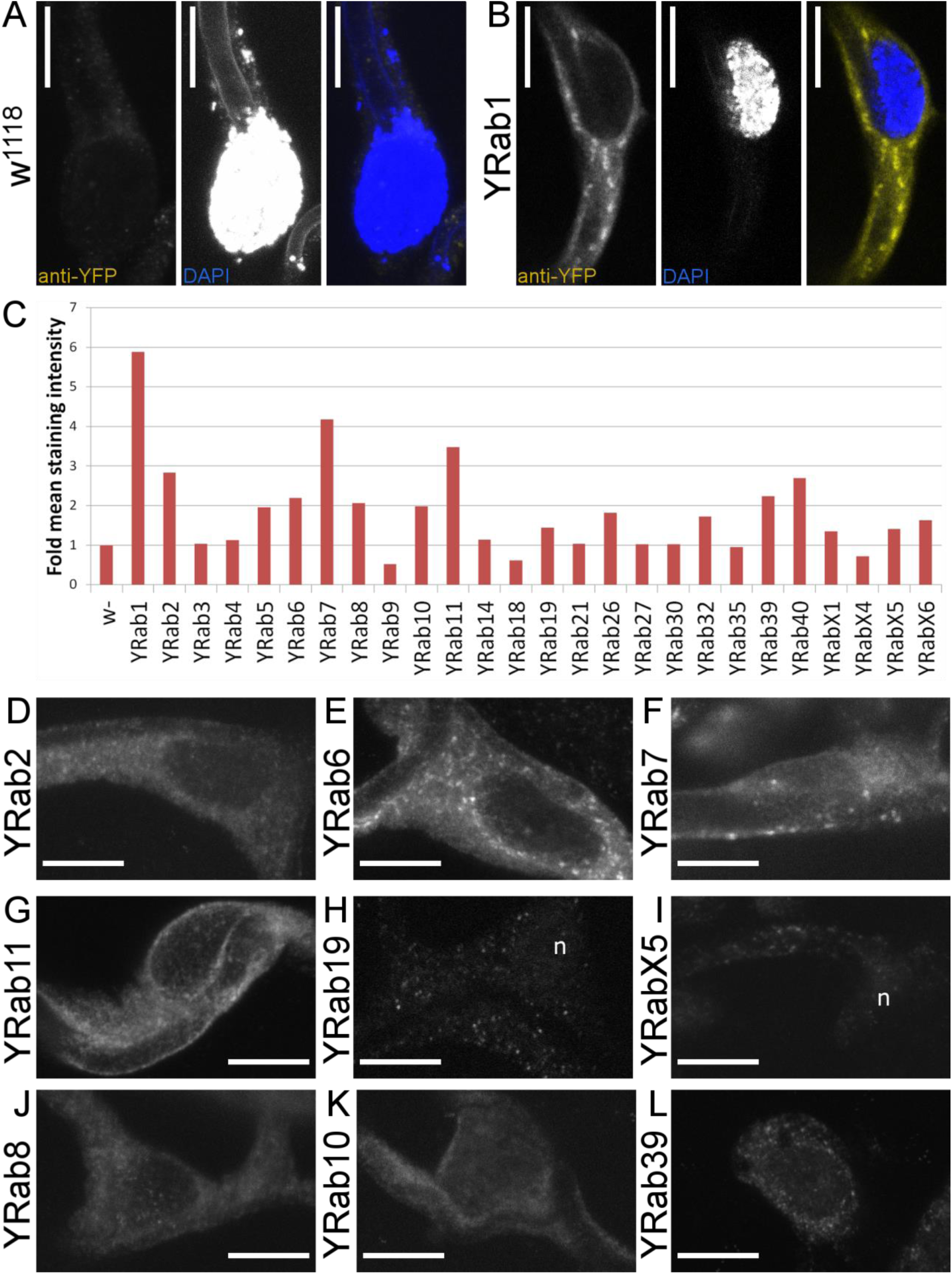
Expression of Rab proteins in Drosophila tracheal terminal cells. L3 wandering stage larvae with endogenously YFP-tagged Rab alleles were dissected and stained against YFP (using an anti-GFP antibody that cross-reacts with the YFP tag). All images show the main body of a dorsal terminal cell (TC) including its nucleus. Scale bars, 10 µm. (A) Negative control TC in a larva expressing no GFP or YFP. All anti-GFP antibodies tested produced a weak punctate staining in negative controls (see Methods). The DAPI channel in this case also captured autofluorescence from the apical extracellular matrix in the subcellular tube’s lumen. (B) TC in a larva with endogenously YFP-tagged Rab1 (YRab1), which shows the strongest endogenous YFP fluorescence and staining out of all YRabs. (C) Quantification of protein expression according to staining intensity relative to negative control. (D-L) Examples of TCs (YFP expression) in larvae of the indicated genotypes. (D-G) Strongly expressed YRabs. These YRabs (1, 2, 6, 7, 11) can be detected both by immunofluorescent staining and by endogenous YFP fluorescence. (H and I) Two YRabs that were undetectable in TCs but nevertheless were associated with strong phenotypes in the knockdown screen (see Fig. 3). n, nucleus. (J-L) Three further YRabs of intermediate expression levels that were associated with TC phenotypes (8 and 10, Fig. 4 and 5) or dorsal branch phenotypes (39, Fig. 2 and 3).

**Figure 2.**
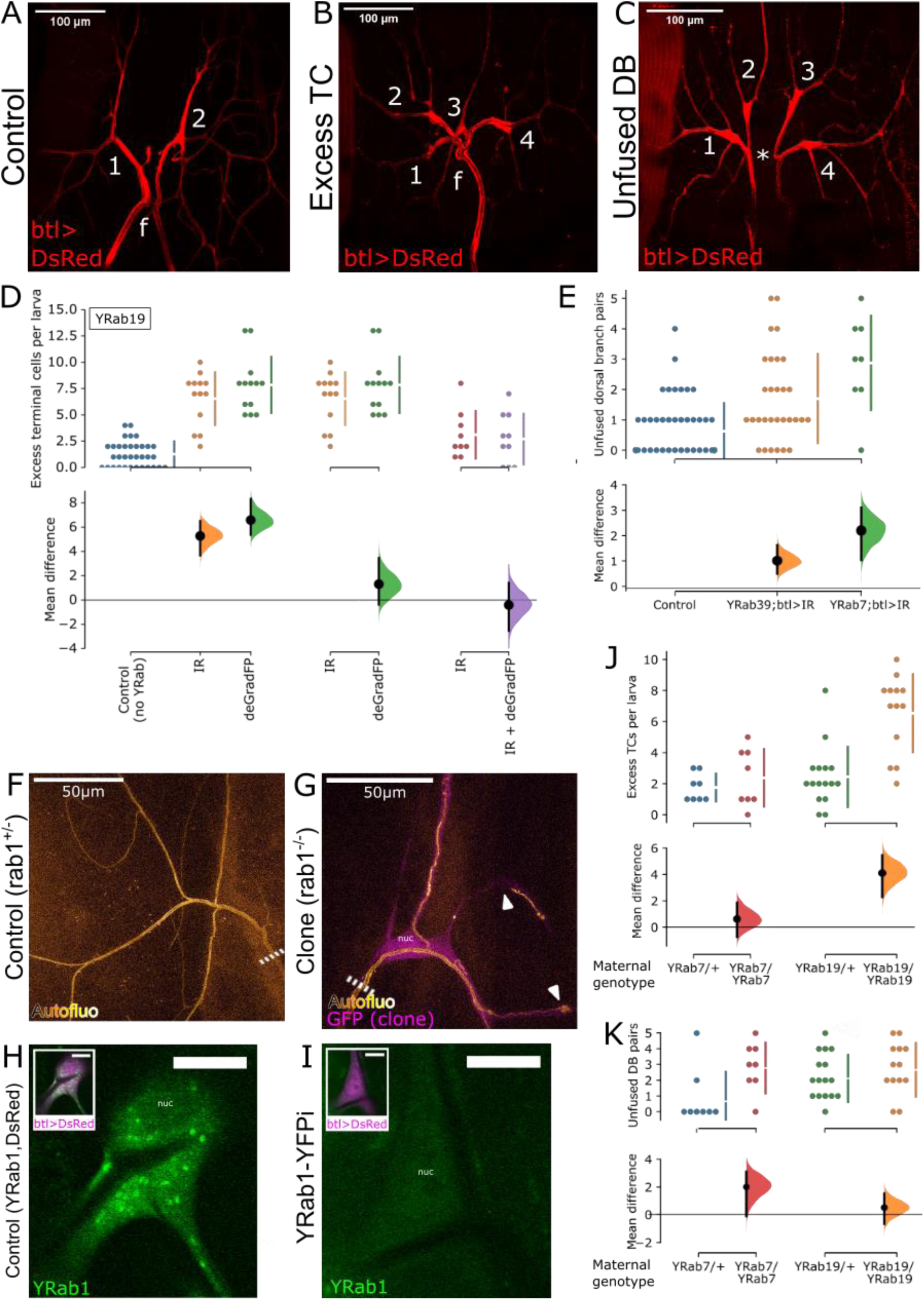
Characterisation of Rab depletion via an endogenously inserted YFP-tag (YRab-YFPi). YRab-homozygous virgins with either a UAS-GFP-knockdown construct or the *btl*-Gal4 and UAS-DsRed constructs were crossed to YRab males with the respective complementary construct(s) to obtain YRab-homozygous larvae expressing the GFP-knockdown transgene and DsRed in tracheal cells. Anterior is up in all micrographs. (A-C) Dorsal branches (DBs) and terminal cells (TCs) of one tracheal metamere in third-instar wandering larvae expressing DsRed and GFP-IR-1 in all tracheal cells. Numbers, terminal cells. f, dorsal fusion. (A) Control larva with no YFP-tagged Rab. Two dorsal branches (DBs) fuse at the dorsal midline. On each DB, one TC ramifies with multiple branches distal to the anastomosis. (B) Example of a tracheal metamere with two excess TCs in a YRab19-YFPi larva. (C) Example of an unfused DB pair (* missing anastomosis), with an excess TC on each side, in a YRab19-YFPi larva. (D) Comparison of two knockdown constructs to deplete YRab19. YRab19-homozygous larvae expressing either GFP-IR-1 (orange, red), deGradFP (green) or both (purple) in tracheal cells were scored for excess TCs. Negative control (blue) refers to larvae expressing GFP-IR-1 under *btl*-Gal4 but with no YRab. The orange and red GFP-IR-1 samples were from separate experiments. Top: dotplots showing number of excess TCs in each larva observed, and mean ± standard deviation (bars next to dotplot). Bottom: estimation of the effect size relative to the respective comparison sample showing mean difference (black dot), 95% confidence interval (black bars) and distribution of bootstrapped mean differences (coloured). (E) Frequency of unfused DB pairs in control (blue), YRab39-(orange) and YRab7-YFPi (green) larvae. Each dot shows the number of unfused DB pairs in one larva (out of in total 8 DB pairs). Estimation statistics are plotted as in (D). (F-G) MARCM clone TCs mutant for *rab1* show abnormalities in the apical extracellular matrix (aECM) that were not reproduced by the YRab1-YFPi knockdown. Compare Fig. 4 and 5. Dashed line, stalk of the TC. (F) Autofluorescence of the aECM in a negative control TC heterozygous for a mutation in *rab1*. This phenotype is identical to wildtype TCs. (G) Homozygous *rab1* mutant TC labelled by cytoplasmic GFP. The aECM reveals abnormalities such as curls near branch tips and disruptions of the tube along stretches of a branch (arrowheads). (H-I) TCs in third-instar wandering larvae with YFP-tagged Rab1 and expressing DsRed in tracheal cells. (H) Control TC not expressing GFP-IR-1 showing strong punctate YFP signal from YRab1. (I) TC expressing GFP-IR-1, showing no detectable YFP signal from YRab1. nuc, nucleus. (J-K) Maternal effect in YRab-YFPi shown by the Excess TC (J) and Unfused DB (K) phenotypes in YRab7- and YRab19-YFPi. Larvae descendent from YRab-heterozygous mothers were compared to larvae descendent from YRab-homozygous mothers. Larvae of the YRab19/YRab19 sample are the same as in (D) since the experiments were carried out in parallel.

**Figure 3.**
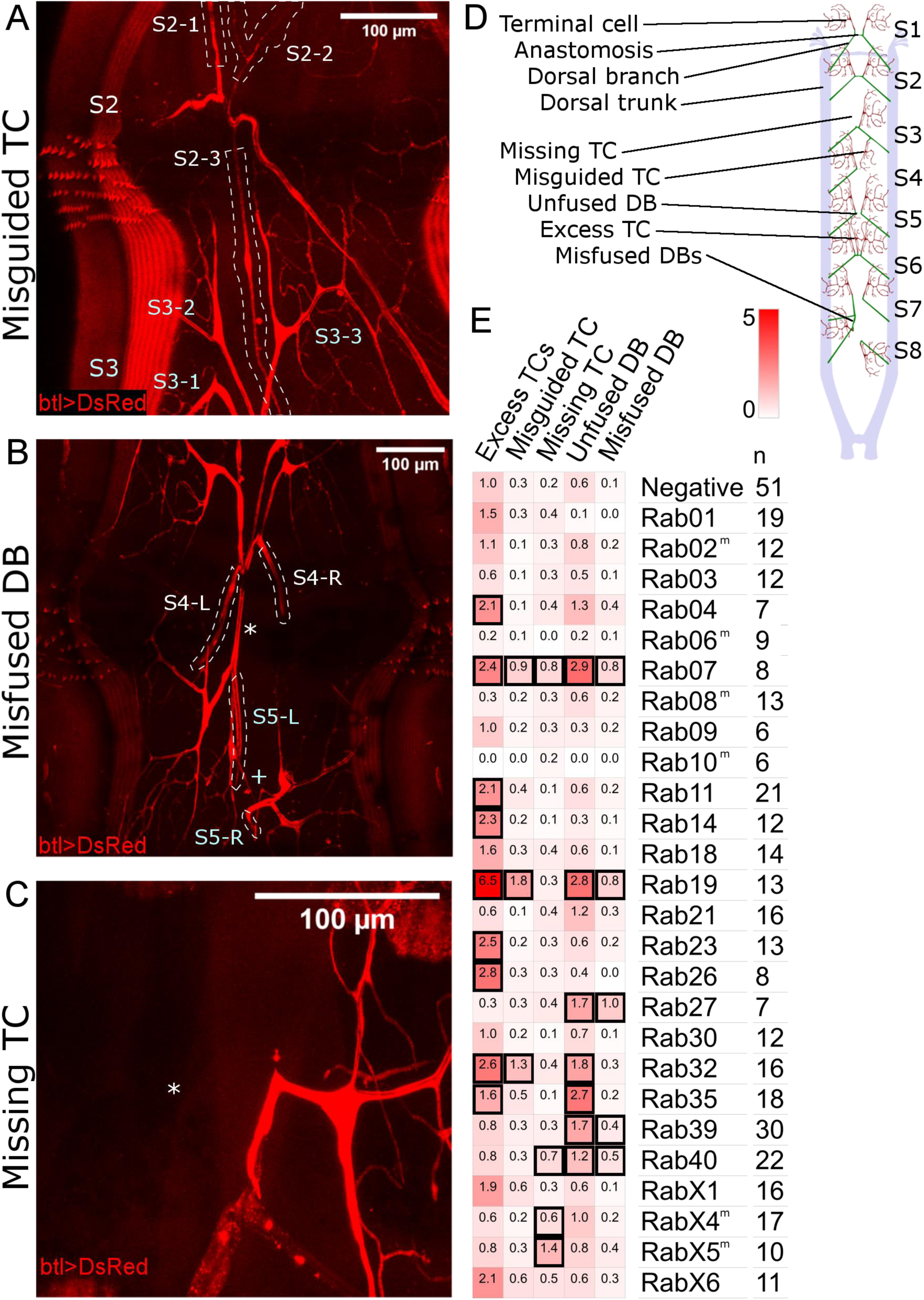
Dorsal branch phenotypes found in the YRab-YFPi screen. YRab-homozygous virgins with either GFP-IR-1, or *btl*-Gal4 and UAS-DsRed were crossed to YRab males with the respective complementary construct(s) to obtain YRab-homozygous larvae expressing the GFP-knockdown transgene and DsRed in tracheal cells. Examples of the “Excess TC” and “Unfused DB” phenotypes can be found in Fig. 2. TC, terminal cell. DB, dorsal branch. Anterior is up in all micrographs. (A) Example of the “Misguided TC” phenotype in a YRab19-YFPi larva. One excess TC (S2-3) on segment 2 (S2) of this larva grew posterior and ramified on a dorsal muscle in segment 3. White dashed lines: outlines of the three TCs of segment 2. (B) Example of the “Misfused DB” phenotype in a YRab19-YFPi larva. The DBs of segment 5 did not fuse in this larva (plus sign). Instead, the left DB of segment 5 (S5-L) formed an anastomosis with a cell of the left DB of segment 4 (S4-L, asterisk). White dotted lines: outlines of the dorsal branches (C) Example of the “Missing TC” phenotype in a YRab9-YFPi larva. The left TC of this larva’s segment 3 is missing (asterisk), though the DBs are fused as normal. (D) Diagram showing the dorsal tracheal anatomy of L3 wandering larvae and the phenotypes observed in the screen. Anterior is to the top. Stereotypically, all DB pairs form a constellation as shown in the first two metameres here. (E) Mean frequencies per larva of each phenotype in YRab-YFPi larvae depleted for the respective Rab. Black squares indicate significant effects (95% confidence interval of difference to control >0).

**Figure 4.**
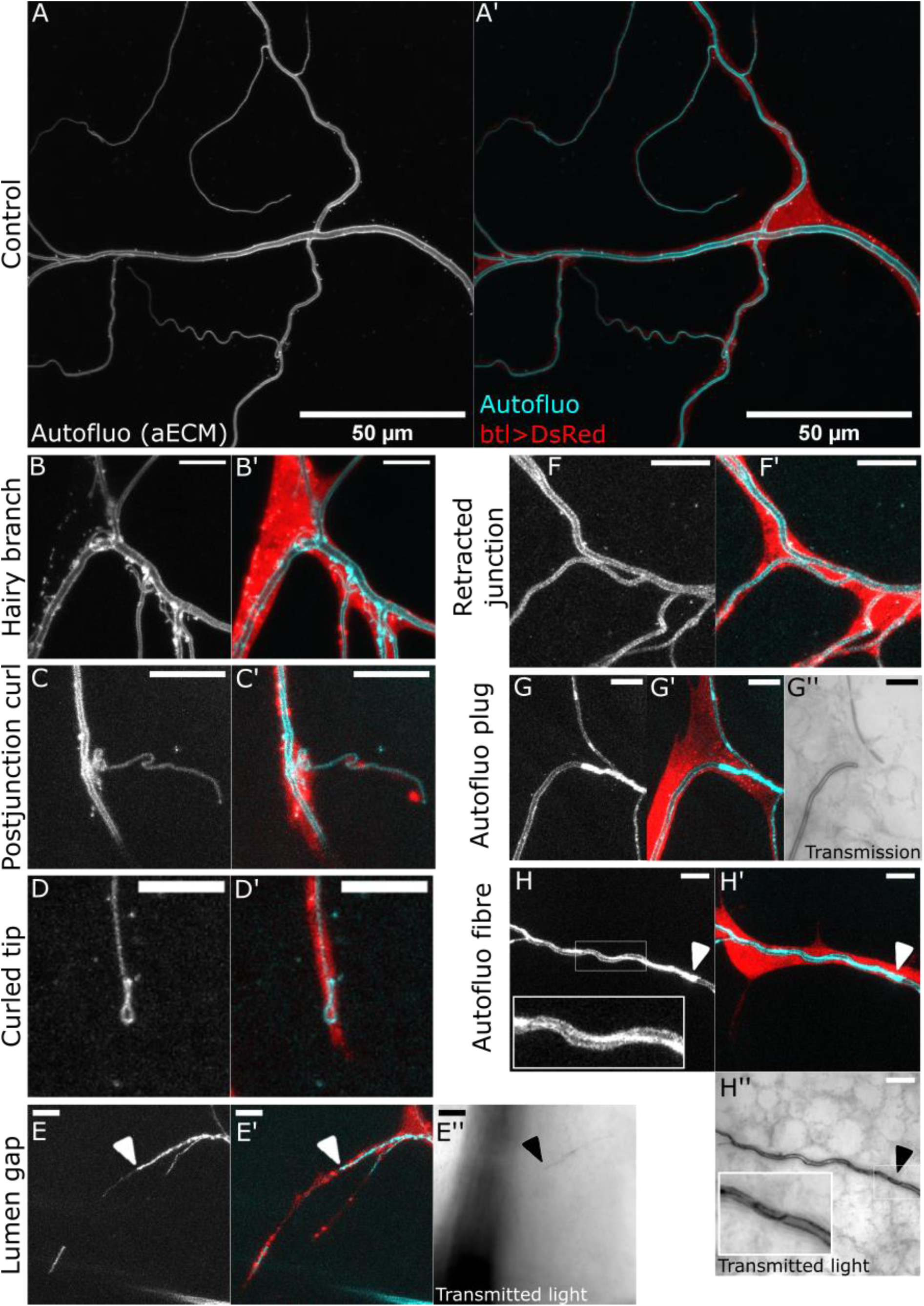
Description of terminal cell phenotypes found in the YRab-YFPi screen. For each case, the first micrograph shows autofluorescence of the apical extracellular matrix (aECM), emitted when illuminated with a 405nm laser. This reveals the morphology of the subcellular tube’s lumen. The second micrograph shows an overlay of autofluorescence (cyan) and cytoplasmic DsRed (red). In some cases, the transmitted light is also shown, which reveals the contrast between the tissue and the gas-filled lumen of the tubes. Scale bars in (A-A’), 50µm; in (B-H”), 10 µ (A-A’) Control dorsal TC in a larva with *btl*-Gal4, GFP-IR-1 and UAS-DsRed. (B-B’) Example of the “Hairy branch” phenotype in a YRab8-YFPi larva. The autofluorescence reveals numerous small tubules and strongly fluorescent puncta surrounding the main subcellular tube. Some of the tubules are connected to the main tube and transmitted light shows that some are gas-filled (not shown). (C-C’) Example of the “Postjunctional curl” phenotype in a YRab8-YFPi larva. The tube is folded up on itself immediately after branching off the parent branch. (D-D’) Example of the “Curled tip” phenotype in a YRab8-YFPi larva. Rather than forming a blunt end, the tube curls back at the branch tip. (E-E”) Example of the “Lumen gap” phenotype in a YRab6-YFPi larva. A stretch of the branch shows no autofluorescence, indicating that no fully mature aECM is present. An aECM-containing tube can be seen distal to the gap. Light transmission (E”) shows that the tube is not gas-filled at or distal to the gap (arrowhead). (F-F’) Example of the “Retracted junction” phenotype in a negative control larva. The points at which the cytoplasm and the tube branch are usually in close proximity. In this phenotype, the point where the tube branches is shifted far away from the point where the cytoplasm branches. (G-G”) Example of the “Autofluorescent plug” phenotype in a YRab6-YFPi larva. A strongly fluorescent “plug” fills the entire diameter of the subcellular tube, with no lumen visible. Gas-filling is also absent in the region of the plug (G”). However, some tubes distal to the plug are gas-filled (see tube at the top in G”). (H-H”) Example of the “Autofluorescent fibre” phenotype in a YRab6-YFPi larva. A strongly fluorescent “fibre” runs along a stretch of the tube (inset in H). (H”) Gas can be seen inside the tube despite the abnormal autofluorescence, although irregularities in the contrast suggest that the autofluorescent fibre protrudes into the lumen (inset in H”).

**Figure 5.**
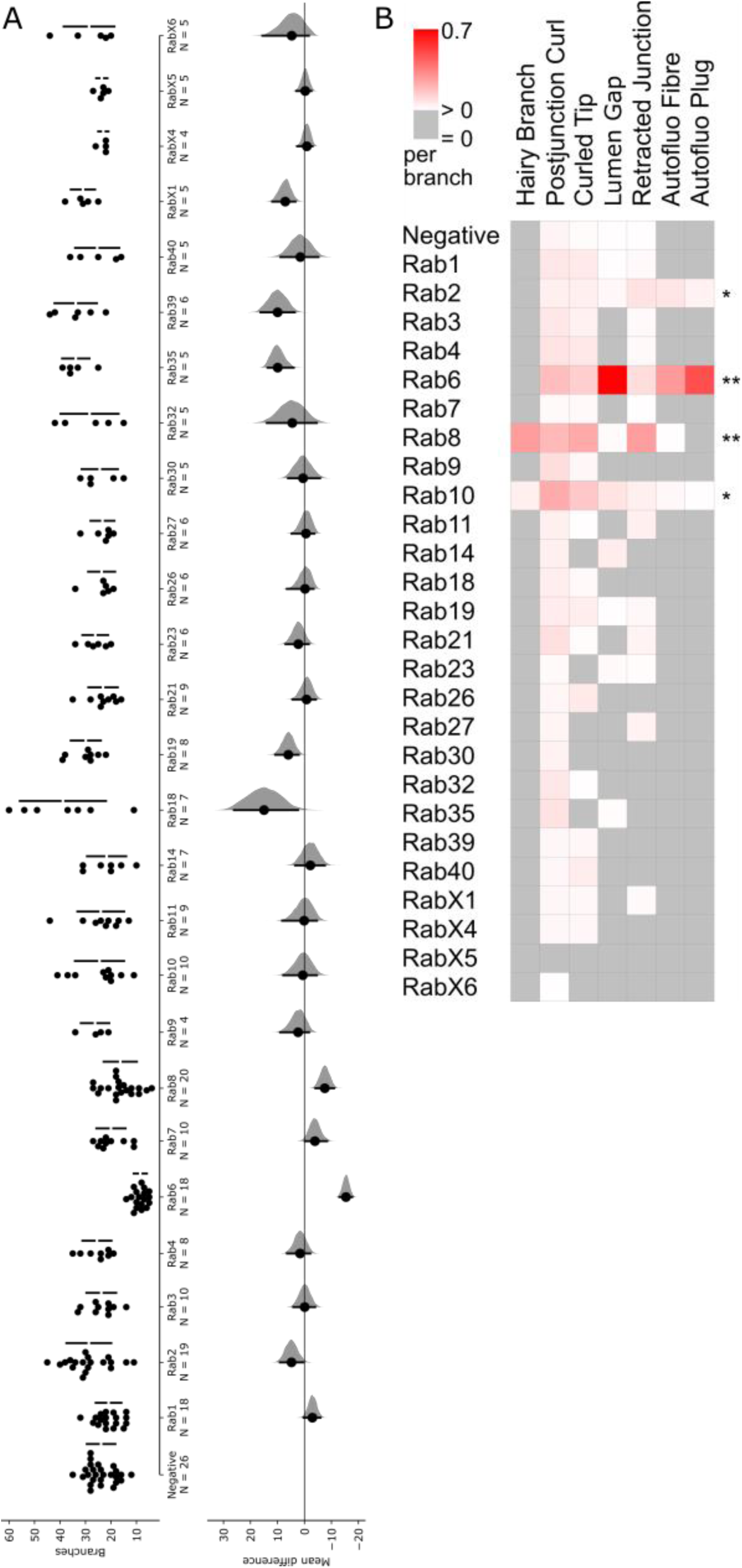
Number of branches and frequency of phenotypes in YRab-YFPi terminal cells. For the initial categorisation of phenotypes, we observed at least 50 dorsal terminal cells (TCs) depleted for each Rab. Once the range of phenotypes had been established, a subset of cells were quantitatively analysed (N for each Rab see panel A). (A) We counted the number of branches per TC depleted for each Rab via YRab-YFPi. This reflects overall cell health (Best, 2019). Left: Dotplots with one dot for each TC and bars showing mean ± standard deviation. Right: Estimation of effect size relative to negative control. Black dot indicates the mean difference to control, black bars indicate the 95% confidence interval, distribution of bootstrapped mean differences in grey. Detailed statistics can be found in Table 1. (B) Mean frequency of phenotypes described in Fig. 4 per branch in the same TCs scored for (A). Asterisks indicate “significant” phenotypes, i.e. those where the autofluorescent aECM reveals differences from control TCs.

### Knockdown of endogenously tagged Rab GTPases by GFP-RNAi or nanobody

The general aim of our screen was to knock down all Rabs of which endogenously YFP-tagged alleles exist, using an indirect approach where the knockdown construct targets the YFP-tag. To identify the most efficient knockdown construct, we first compared the ability of different RNA interference (RNAi) and nanobody constructs that were originally designed against GFP to eliminate YFP fluorescence of tagged Rab1, because this Rab had the strongest endogenous YFP signal. This narrowed our choice down to one RNAi and one nanobody construct, which we then compared in more detail by their ability to elicit an easily quantifiable loss-of-function phenotype related to Rab19 in the tracheal system. We finally chose the RNAi construct for all further work. We validated the indirect knockdown approach by testing its recapitulation of published tracheal phenotypes related to Rabs, and explored the capability of this approach to identify maternal contribution.

#### Selection of knockdown construct for YRab-YFPi

The principle of knocking down an endogenously tagged protein by targeting the tag has been demonstrated in several previous studies. We compared the knockdown efficiency of two GFP-RNAi constructs (GFP-IR1 from NIG and EGFP.shRNA.3 from BDSC) and two different nanobody insertions (deGradFP either in a P-element or an M-element) by imaging endogenously YFP-tagged Rab1 (YRab1). YRab1 is expressed at high levels in the larval epidermis, which we used for this optimisation because it is easy to image and the genetic tools allow obtaining an internal control within one animal. Due to the structural similarity of GFP and YFP, most constructs recognise both tags. Each knockdown construct was expressed under *engrailed-gal4* to obtain epidermal cells expressing the construct juxtaposed to cells not expressing it, and the level of YFP fluorescence in the neighbouring cell groups was compared to measure knockdown efficiency. We found that out of the four knockdown constructs, GFP-IR1 eliminates the YFP signal with the highest efficiency (Fig. S1), followed by the other RNAi construct. Nanobody-mediated knockdown reduced YFP signal but sometimes failed to eliminate the punctate structures characteristic for YRab1. We also verified that GFP-IR1 depletes YFP fluorescence of YRab1 in tracheal terminal cells when driven by *breathless*-gal4 (*btl-gal4*) (Fig. 2H-I).

#### Comparison of RNAi and deGradFP at phenotype level

We considered the possibility that the fluorescence of the YRab1 protein might not necessarily correspond to its Rab activity, for example if the nanobody was able to inactivate YRab1 protein by binding it [find ref for antibody binding can modify structure of protein], but did not successfully target it for degradation. To test this, we compared the phenotype severity elicited by GFP-IR-1 to deGradFP using an easily quantifiable defect. We had noticed that in larvae where Rab19 is depleted under the *btl-gal4* driver, dorsal branches frequently had excess TCs (Fig. 2B, D). When we compared the effect of GFP-IR-1 to deGradFP, we found that the frequency of excess TCs was similar when YRab19 was knocked down by GFP-IR-1 compared to deGradFP, or compared to the combination of both (Fig. 2D). Thus, even though deGradFP did not eliminate YRab1 fluorescence fully in the previous test, it achieves a similar level of Rab19 loss-of-function as GFP-IR-1, and combining both did not further improve the knockdown efficiency. We therefore suggest that deGradFP may be capable of interfering with the function of other protein domains simply by binding the fluorescent tag domain, even without targeting the entire protein for degradation. To further confirm that GFP-IR-1 and deGradFP are functionally equivalent knockdown approaches, we tested both against the preliminary “Rabs of interest” for terminal cells, i.e. those with the strongest expression level (Rab1, 7 and 11), and Rab8 due to its striking loss-of-function phenotype in terminal cells (described later). For Rab1, 7 and 11, neither of the two knockdown approaches resulted in phenotypic abnormalities. For Rab8, the phenotypic defects were of indistinguishable severity with both approaches (cf. Fig. 5 showing the results for RNAi). We therefore conclude that the loss-of-function achieved by tag-mediated knockdown approaches is generally equivalent regardless of whether the mRNA or protein product is targeted, and combining both approaches does not achieve a higher knockdown efficiency. For our knockdown screen, we chose to proceed using GFP-IR-1.

#### Consistency of YRab-YFPi with Rab-related tracheal phenotypes

To test the validity of YRab-YFPi, we compared the knockdown phenotypes to those that have already been published for Rab7, Rab10 and Rab11, Rab35 and Rab39. We could not apply this to Rab5, because even minimal knockdown of YRab5 was lethal for larvae. Thus, no viable YRab5-homozygous larvae hatched, regardless of whether we used *btl-gal4* or *dsrf*-*gal4*, which is expressed in TCs and muscles. YRab5-heterozygous larvae showed no abnormalities. In YRab7- and YRab39-YFPi larvae, we observed an increased frequency of unfused dorsal branch pairs (Fig. 2C & E), consistent with a requirement for Rab7 and Rab39 in this fusion process (Caviglia et al., 2016). Rab10-depleted TCs sometimes (Fig. 5B) showed a defect in the apical extracellular matrix that we termed “autofluorescent plug” (Fig. 4G), consistent with the tube collapse reported for TCs overexpressing an inactive Rab10 mutant variant (Jones et al., 2014). We identified no abnormal phenotypes in Rab11- or Rab35-depleted TCs, and none in Rab11-depleted DBs that would correspond to the role of Rab11 in maintaining adherens junctions (Le Droguen et al., 2015). One possible explanation for the discrepancy between our results and the reported phenotypes is that the latter resulted from overexpressing inactive Rab mutant variants. Thus, either the YRab-YFPi in our experiments did not deplete the Rabs sufficiently, or the reported phenotypes reflect ambiguous-dominant effects rather than loss of function. The overexpressed mutant Rabs could for example not only interfere with their wildtype counterpart’s function, but simultaneously disrupt the function of other Rabs on the same membrane compartment.

**Table 1.** Branch counts and statistics estimating impact of YRab-YFPi on terminal cell branch number.

| Tagged gene | Mean TC branch count | Branch count std dev | Mean difference to control | 95% CI lower bound | 95% CI upper bound | CI threshold | p-value | Sign. |
| --- | --- | --- | --- | --- | --- | --- | --- | --- |
| Negative | 23.9 | 5.6 |  |  |  |  |  |  |
| Rab1 | 20.9 | 5.0 | -2.9 | -5.9 | 0.3 |  | 0.06 |  |
| Rab2 | 28.7 | 8.8 | 4.8 | 0.2 | 9.1 | increase | 0.017 | * |
| Rab3 | 23.8 | 5.8 | -0.1 | -3.9 | 4.1 |  | 0.94 |  |
| Rab4 | 25.5 | 5.7 | 1.6 | -2.1 | 6.5 |  | 0.58 |  |
| Rab6 | 8.5 | 2.6 | -15.4 | -17.7 | -12.9 | decrease | 2.90E-08 | *** |
| Rab7 | 20.0 | 5.7 | -3.9 | -8.4 | -0.1 | decrease | 0.07 |  |
| Rab8 | 16.4 | 6.5 | -7.5 | -11.0 | -4.1 | decrease | 2.55E-04 | *** |
| Rab9 | 26.3 | 5.6 | 2.4 | -1.7 | 9.0 |  | 0.65 |  |
| Rab10 | 24.5 | 9.6 | 0.6 | -4.6 | 7.9 |  | 0.83 |  |
| Rab11 | 24.0 | 9.2 | 0.1 | -4.6 | 8.3 |  | 0.61 |  |
| Rab14 | 21.7 | 7.7 | -2.2 | -7.6 | 3.4 |  | 0.63 |  |
| Rab18 | 38.9 | 16.9 | 15.0 | 2.4 | 26.1 | increase | 0.0117 | * |
| Rab19 | 29.9 | 5.9 | 6.0 | 2.1 | 10.8 | increase | 0.025 | * |
| Rab21 | 23.1 | 5.8 | -0.8 | -4.2 | 4.4 |  | 0.53 |  |
| Rab23 | 26.2 | 5.0 | 2.3 | -1.6 | 7.0 |  | 0.44 |  |
| Rab26 | 23.7 | 5.3 | -0.2 | -3.4 | 6.5 |  | 0.68 |  |
| Rab27 | 23.3 | 4.7 | -0.6 | -3.7 | 4.8 |  | 0.81 |  |
| Rab30 | 24.4 | 7.1 | 0.5 | -5.8 | 6.1 |  | 0.73 |  |
| Rab32 | 28.4 | 11.4 | 4.5 | -4.5 | 13.9 |  | 0.55 |  |
| Rab35 | 33.8 | 5.4 | 9.9 | 4.0 | 13.6 | increase | 0.006 | ** |
| Rab39 | 33.8 | 8.3 | 9.9 | 3.6 | 16.3 | increase | 0.012 | * |
| Rab40 | 25.4 | 8.6 | 1.5 | -5.2 | 8.9 |  | 0.77 |  |
| RabX1 | 31.0 | 4.7 | 7.1 | 3.4 | 11.9 | increase | 0.013 | * |
| RabX4 | 23.0 | 2.0 | -0.9 | -3.0 | 2.7 |  | 0.54 |  |
| RabX5 | 23.6 | 2.2 | -0.3 | -2.6 | 3.0 |  | 0.63 |  |
| RabX6 | 28.6 | 9.9 | 4.7 | -1.6 | 15.6 |  | 0.43 |  |

P-values according to Mann-Whitney U test. CI, 95% confidence interval of the bootstrapped distribution of mean differences. Std dev, standard deviation. Sign., significance.

#### Degree of loss-of-function caused by indirect RNAi with btl-gal4

Errors in knockdown screens can arise from false positives (e.g. when the knockdown construct affects other genes than the one targeted), and false negatives (e.g. when the knockdown does not efficiently eliminate the gene product). The YRab-YFPi method provides a strong control against false positives, because the tag-knockdown construct can be expressed in the absence of a tagged target. However, we suspected that false negatives may be present because some phenotypes associated with perturbations of Rabs were not reproduced by YRab-YFPi in the previous experiments. We also generated TCs with a null mutation in *rab1* (*rab1*^*S147213*^; Sechi et al., 2017), using MARCM (Lee and Luo, 2001) and compared their phenotype to TCs in YRab1-YFPi larvae. Mutant TCs had severely reduced branch numbers and/or abnormalities in their apical extracellular matrix (aECM; n=6) (Fig. 2F-G). In contrast, TCs depleted of YRab1 had no aECM abnormalities and normal branch numbers (Fig. 5). Yet, YRab1-YFPi completely abolished the fluorescent signal from YRab1 (Fig. 2H-I). We therefore conclude that the loss-of-function caused by YRab-YFPi for Rab1 is either incomplete, and an undetectable amount of residual YRab1 protein is sufficient for TCs to undergo normal morphogenesis, or else, that the chromosome carrying the Rab1 mutation also has other mutations on it that affect tracheal branching morphogenesis.

#### Maternal contributions of Rab7 and Rab19

A further concern of genetic interference with gene function is maternal contribution. Since RNAi only eliminates mRNA, maternally deposited protein products of the targeted gene are not eliminated. Depending on the protein’s stability, it can be sufficient to partially or completely execute the gene’s function, and thus mask loss-of-function phenotypes. The YRab-YFPi method makes investigating the impact of maternal contribution simple, because only YFP-tagged gene products are knocked down. If there is a maternal contribution for any given Rab, mothers heterozygous for the YRab allele deposit both tagged and untagged gene products. The untagged product cannot be knocked down in the progeny and would provide some of the gene’s function during development. By contrast, in embryos derived from YRab-homozygous mothers, all of the gene product is tagged and therefore targeted by YRab-YFPi. Loss-of-function phenotypes should thus be more severe in progeny of YRab-homozygous mothers, if the maternally deposited product participates in the corresponding developmental process. We noticed a divergence of phenotypes in preliminary YRab-YFPI crosses for Rab7 and Rab19 and therefore tested the dependence on maternal genotype for these. For both YRabs, knockdown larvae had excess TCs on their dorsal branches (Fig. 2B), and some dorsal branch pairs were not fused as is normally the case (Fig. 2C). In case of YRab7, the number of excess TCs was not dependent on the genotype of the mother (Fig. 2J), but the number of unfused dorsal branches was (Fig. 2K). For YRab19 the reverse was the case, the frequency of excess TCs was strongly affected by the maternal contribution (Fig. 2J), but the number of unfused dorsal branches was not (Fig. 2K). Maternally deposited *rab7* and *rab19* gene products therefore contribute to dorsal branch morphogenesis. Since different phenotypes were affected by maternal genotype, it is likely that each Rab’s gene dosage is limiting for different developmental processes. We tested YRab11 and YRab32 for maternal contribution in the same way and similarly observed lower phenotypic penetrance in larvae from YRab-heterozygous mothers.

### Rabs required for cell fate specification and fusion in dorsal branches

To screen for knockdown phenotypes, we focussed on the dorsal part of the larval tracheal system at L3 wandering stage (Fig. 3D). We found two categories of defects: those relating to the morphogenesis of the tracheal dorsal branch (DB), characterised by incorrect numbers of terminal cells (TCs) or failed connections of fusion cells across the dorsal midline (this section), and those relating to the subcellular morphogenesis of dorsal TCs, including incorrect numbers of branches and more subtle defects in cell shape (next section).

In the first category, we found five types of abnormalities:

- excess TCs (Fig. 2B), where the additional TC’s stalk was attached either to the DB proximal to the fusion point, on the fusion bridge itself, or on another TC’s stalk distal to the fusion point;
- unfused DB pairs (Fig. 2C);
- misguided TCs (Fig. 3A), growing either posterior to the fusion point into the next segment, anterior into the previous segment, laterally or interior, usually failing to tracheate the stereotypical target muscle;
- misfused DBs (Fig. 3B), where fusion occurred laterally with the DB from the previous or next segment, or diagonally across the midline with a DB from another segment - we note that misfused DBs in some cases fused both with a partner from a neighbouring segment, but also with a contralateral DB. This suggests that the fusion cells of DBs have the capacity to fuse with more than one partner fusion cell;
- and lastly missing TCs (Fig. 3C).

Note that all five phenotypes were sometimes found in negative control larvae (expressing GFP-RNAi but without tagged Rab) and in wildtype larvae (Oregon-R strain). Thus, the phenotypes themselves cannot strictly be considered “defects”, but would be better described as deviations from the stereotypical configuration of cells at DB tips. Stereotypically, eight of the eleven larval body segments each contain two bilateral DBs that are fused at the dorsal midline (Fig. 3D). Distal to this fusion point, one TC ramifies on each side (Fig. 2A).

We quantified the frequency of each abnormality in YRab-YFPi larvae (Fig. 3E). We found 14 Rabs that were associated with significant increases in the frequency of at least one abnormality over control (i.e. 95% confidence interval of the difference between knockdown and control was larger than 0). Five Rabs were associated only with increased frequencies of excess TCs (Rab4, 11, 14, 23, 26). Two Rabs were associated only with missing TCs (RabX4, X5). Two Rabs were associated with both unfused DBs and misfused DBs (Rab27, Rab39). The remaining five Rabs each showed a unique combination of defects (Rab7, 19, 32, 35, 40). Rab7 was the only Rab associated with increased frequencies of all five types of defects. Most of these phenotypic profiles can be explained by mistrafficking of signalling molecules, which we discuss below.

### Rabs required for terminal cell morphogenesis

To assess TC morphogenesis, we scored the number of branches per TC as the primary outcome phenotype (Fig. 5A). Only two YRab-YFPi samples showed strongly reduced branch counts: Rab6 and Rab8. Rab7 was associated with a small but statistically insignificant reduction of branch counts. Rab2, 18, 19, 35, 39, X1 were associated with significantly increased average branch counts. The effects on branch numbers were slight (though significant) in most cases, and only Rab18 knockdown resulted in a more dramatic increase in the numbers of branches that went well outside of the range we saw in controls.

Virtually every cellular pathway that has been studied in TCs, when perturbed, is associated with changes in the number of branches (Best, 2019). We therefore consider branch number an unspecific phenotype that reflects overall TC “health”. We thus looked for qualitative differences in the morphology of the cells’ cytoplasm and their subcellular tube network. We used the autofluorescent apical extracellular matrix to observe subcellular tube morphology (Fig. 4A). We first screened at least 50 TCs for each YRab-YFPi sample to compile a list of morphological abnormalities (Fig. 4). We then quantified the frequency of all of these features in at least 5 TCs for each YRab-YFPi sample. Their frequencies per TC branch are shown in Fig. 5B. We found no abnormalities relating to the shape of the cytoplasm (i.e. the cell soma or branches).

Four Rabs were associated with aECM defects that were never found in controls, or increased frequencies of phenotypes that also appear in controls: Rab2, Rab6, Rab8 and Rab10. The most striking phenotypes were associated with Rab6 and Rab8. Rab6-depleted TCs showed frequent disruptions of the aECM and gas-filling (“gap”, Fig. 4E), indicating a collapse of the tube, as well as aggregations of autofluorescent material on one side of the tube resembling a long fibre (“autofluorescent fibre”; Fig. 4H). In some cases, an aggregate of autofluorescent material occupied the entire tube diameter like a “plug”, correlating with a lack of gas-filling (Fig. 4E). Rab2-depleted TCs showed the same abnormalities, although the severity and frequency was markedly lower (Fig. 5B), and branch numbers were unaffected by YRab2-YFPi (Fig. 5A).

Rab8-depleted TCs had defects in the number and branching of tubes. In some branches, the main tube was enveloped with numerous thin tubules branching off from it (“hairy branch”, Fig. 4B). Some of these tubules contained a mature aECM as shown by the presence of taenidia and gas-filling, while others were not connected to the main tube and had no visible lumen of their own. Rab10-depleted TCs, although rarely, showed similar “specks” on the main tube as also seen in Rab8-depleted TCs with fewer “hairs”. We therefore interpret this as a particularly mild form of the “hairy branch” phenotype and have scored it accordingly for the quantification (Fig. 5B).

Similar to previously reported phenotypes of tube bundling at branch tips (Levi et al., 2006; Schottenfeld-Roames and Ghabrial, 2012), we also found morphologies where the tube tip was not straight (“curled tip”) (Fig. 4D). These were present at a low frequency in control cells (Fig. 5B) and at a higher frequency in YRab-YFPi against Rab6, Rab8 and Rab10. With Rab8 knockdown, curled tips were also often larger than typically found in controls. We further noticed instances of the tube folding after branching (“postjunctional curl”, Fig. 4C), as well as instances where the branching point of the tube was located far away from the cytoplasmic branching point (“retracted junction”, Fig. 4F). Both phenotypes were sometimes found in control cells but were more frequent in Rab6, Rab8 and Rab10 knockdown (Fig. 5B).

## Discussion

### Implications of *btl-gal4* expression pattern for YRab-YFPi

The *btl-gal4* driver has been used to drive transgene expression specifically in tracheal cells for decades (Shiga et al., 1996). Throughout our experiments, we consistently saw expression of DsRed in some epidermal cells when driven by *btl-gal4*, whereas the anterior and posterior anastomoses between the dorsal trunks contained tracheal cells that never expressed DsRed (data not shown). These details of the spatial expression pattern of *btl-gal4* are unlikely to affect any of the processes we studied here. However, the expression of *btl-gal4* across time needs to be considered when interpreting our screen results. Transcription of the *btl* gene itself peaks during mid-embryogenesis, coinciding with the onset of tracheal pit invagination (Brown et al., 2014). It then declines rapidly until the end of embryogenesis and remains at low but detectable levels throughout larval life. The expression of *btl-gal4*, the induction of UAS-controlled transgenes, and the silencing induced by RNAi expressed this way, do not necessarily follow the same kinetics as the expression of *btl* mRNA. There are currently no data available to infer whether or not they do. However, the strong *btl* expression peak during the early stages of tracheal development could cause YRab-YFPi to have a high efficiency during the period when the tracheal primary and dorsal branches are forming. Later, when the terminal cells (TCs) ramify during the larval stages, the lower expression of *btl* could mean that also the RNAi construct is expressed at weaker levels. This could be one factor contributing to the high discovery rate of Rabs relevant for dorsal branch morphogenesis - which occurs during the embryonic stages - while very few Rabs were identified as relevant for terminal cell morphogenesis occurring later in development.

### Relation between Rab expression level and knockdown efficiency

Regardless of the efficiency of YRab-YFPi across space and time, its expression should be comparable in all YRab-YFPi experiments. Therefore, all YRabs in our screen were subject to the same perturbation. However, the Rabs themselves were expressed at different levels in TCs, and one could expect this to cause differences in the knockdown efficiency and therefore the likelihood of discovering loss-of-function phenotypes. The Rabs with the highest expression levels in larval TCs were YRab1, 6 and 11. For all three, no residual signal from the YFP-tag was detectable in TCs in the YRab-YFPi knockdown condition. Yet, neither Rab1 nor Rab11 depletion caused any defects in TC morphogenesis, while Rab6 was associated with the most penetrant loss-of-function phenotype in our screen. At least for Rab1, the morphological defects in *rab1* mutant TCs suggested that the protein does play an important role in their morphogenesis. Unless this is a false positive due to confounding mutations on the *rab1* mutant chromosome, this would suggest that YRab-YFPi does not eliminate all YRab1 protein. In contrast, many YRabs had barely detectable expression in TCs, but out of these, only Rab8 depletion caused morphological defects. For YRab19, combining RNAi and protein degradation via nanobody did not cause more penetrant dorsal branch abnormalities than either knockdown alone. Thus, the same knockdown that may have been incomplete for YRab1, may have reached saturation for YRab19. The quantitative connections between RNAi efficiency, Rab expression and phenotype severity therefore cannot simply be generalised. This is probably due to the unique functions of each Rab, and differences in the knockdown efficiency that would be required to elicit any defects in morphogenesis.

### Involvement of Rabs in dorsal branch morphogenesis

The YRab-gfpIR screen revealed a large number of Rabs as required for proper morphogenesis of the dorsal branch (DB). Among these are as yet uncharacterised genes such as RabX5 and Rab19, but also well-known members of the family such as Rab7 and Rab11. The developmental processes that underlie the DB defects observed here are sufficiently well understood to suggest causative mechanisms, all of which take place during embryogenesis. Given that Rabs are involved in membrane trafficking, most cases can be explained by assuming that the Rab is needed to deliver a receptor involved in cell fate specification or migration to the appropriate membrane compartment. In this section, we interpret the knockdown phenotype profile associated with each Rab to predict the signalling receptors that require the Rab.

Cell fate determination within the DB can be subdivided into three steps (Hayashi and Kondo, 2018):

1. Specification of DB cells from the tracheal placode in response to Decapentaplegic (Dpp) signalling from the dorsal midline (Chen et al., 1998; Vincent et al., 1997), followed by DB “sprout” formation. This step potentially occurs before the expression onset of transgenes driven by *btl-gal4* and may therefore be unaffected by the knockdown.
2. Selection of tip cells within the DB group, followed by migration of the tip cells and intercalation of the remaining cells to form the DB stalk. During this competitive process, cells with higher Btl activity migrate to the tip of the group (Du et al., 2017; Kato et al., 2004; Le Droguen et al., 2015), and lateral inhibition mediated by Delta and Notch restricts the number of tip cells to two (Ohshiro et al., 2002; Schottenfeld et al., 2010).
3. Specification of tip cells into fusion cells and terminal cells, followed by individual cellular morphogenesis. The cell closest to the dorsal source of Wingless (Wg) and possibly other Wnts, as well as Dpp adopts the fusion cell fate (Llimargas, 2000; Steneberg et al., 1999). This leads it to stop expressing Btl and upregulate Delta expression instead. (Chihara and Hayashi, 2000). The neighbour cells’ resulting Notch activation suppresses the fusion cell fate. The other tip cell thus continues expressing Btl, and adopts the terminal cell fate (Guillemin et al., 1996).

The most prevalent abnormality was an increased number of terminal cells (TCs). This defect could arise either during selection of two tip cells from the DB group or during fate refinement of tip cells into TC and FC. Five Rab knockdowns (Rab4, Rab11, Rab14, Rab23 and Rab26) caused excess TCs but not unfused DBs (i.e. FCs were present and functional), indicating a net increase in tip cells. Since the number of tip cells is restricted by Notch activity (Ikeya and Hayashi, 1999; Llimargas, 1999), these mild phenotypes could reflect lowered Notch signalling.

Indeed, Rab4 is involved in Notch trafficking in wing discs (Gomez-Lamarca et al., 2015) and the human homologue Rab4a recycles Notch from endosomes to the plasma membrane in HeLa cells (Zheng and Conner, 2018). Rab11 deficiency is associated with impaired secretion of Delta in sensory bristle development (Charng et al., 2014), which implies lower Notch activation. Rab14, 23 and 26 have so far not been associated with Notch signalling.

A defect where the number of tip cells is normal but both adopt the TC fate (instead of one TC and one FC), manifests phenotypically as an unfused DB with an excess TC. In addition, if both tip cells become TCs, the excess TC is more likely to mismigrate (Kato et al., 2004). This combination of excess TC, unfused DB and misguided TC occurred in Rab7, Rab19 and Rab32 knockdown. Kato and colleagues observed this when Wg signalling was inhibited. Alternatively, excess activity of Btl can also lead to the specification of excess TCs at the expense of fusion cells (Ghabrial and Krasnow, 2006; Lebreton and Casanova, 2016; Sutherland et al., 1996). This suggests that Rab7, 19 and 32 are required for trafficking Wg receptors, and/or for inactivating or degrading active Btl.

Missing TCs were overall very rare, but the two Rabs for which this phenotype was characteristic, RabX4 and RabX5, have barely been characterised so far. Selecting only a single tip cell out of the DB group would suggest excess Notch signalling, but at the later transition of tip cells into TCs and FCs, excess Notch represses the FC fate (Araujo and Casanova, 2011; Ikeya and Hayashi, 1999) and should cause a single tip cell to become a TC rather than a FC. We therefore speculate that RabX4 and RabX5 are involved in Notch trafficking only during the early stages of tracheal development.

The Unfused DB phenotype could result from an absent fusion cell, or from defects in the fusion process itself. Rab7 is implicated in the fusion process (Caviglia et al., 2016), although the presence of other DB abnormalities in Rab7 knockdown indicates that this is not its only function in DB morphogenesis. Rab27, Rab39 and Rab40 knockdown caused unfused DBs, without concomittant excess TCs. The normal number of TCs is a likely indicator of a defect in the fusion process rather than fate specification. Rab39 is already known to be involved in secretion into the newly forming fused lumen during this process (Caviglia et al., 2016). The secretion pathway in this fusion process proceeds through lysosome-related organelles, a mechanism that involves Rab27 in immune cells (Fukuda, 2013). Furthermore, one Rab27 effector regulating lysosome secretion in haematopoietic cells (Neeft et al., 2005) and platelets (Shirakawa et al., 2004) is Munc-13-4, the homologue of Staccato, which was identified along with Rab39 on secretory lysosomes in tracheal fusion cells (Caviglia et al., 2016). Rab27 is therefore a strong candidate for contributing to fusion along with Rab39. Rab40 was also associated with Missing TCs and therefore probably participates in other processes besides fusion.

Finally, the abnormality Misfused DB most likely involves a migratory defect, in which the FC transgresses the boundary of the body segment. This suggests an insensitivity to the guidance probably provided by the epidermal cells along which tracheal tip cells migrate (Kato et al., 2004; Le Droguen et al., 2015). The respective Rabs (Rab7, Rab19, Rab27, Rab39, Rab40) could thus be involved in trafficking a membrane-bound signalling molecule whose function involves direct cell-to-cell contact, such as integrins. However, which receptors mediate the contact between the DB and the muscle or epidermis is not known.

### Involvement of Rabs in terminal cell morphogenesis

To identify Rabs with important roles in terminal cell (TC) morphogenesis, we first scored the number of branches formed by TCs to obtain an unspecific readout of TC “health”. Second, we looked for a method of assessing TC morphology that could provide information on the molecular pathway in which a Rab might be involved. During a preliminary screening, we saw that control TCs frequently had branches whose tubes appeared “abnormal”, for example with a curled tube instead of a straight one. We also observed that even TCs with striking defects often still formed a subset of branches with normal morphology. This led us to a quantitative approach, scoring the frequency of morphological abnormalities/features per branch.

Those Rabs whose depletion caused strong reductions in branch numbers (Rab6 and Rab8) were also associated with the most severe phenotypes in their subcellular tubes, consistent with our previous conclusion that TCs’ ability to form branches depends on their ability to form proper tubes (reviewed in Best, 2019). Rab2 and Rab10 showed abnormal tube features, albeit at low enough frequencies that the cells’ ability to form branches was not affected. In fact, Rab2-depleted TCs had more branches than control TCs. Phenotypes with increased TC branch numbers have so far only been associated with FGF and insulin signalling (Jones and Metzstein, 2011; Wong et al., 2014). Rab2 may thus be involved in the trafficking of the FGF receptor, Btl, or the insulin receptor.

Two of the three morphological features that we observed in Rab2- and Rab6-depleted TCs, autofluorescent “plugs” and lumen gaps, can be explained by mechanical instability of the aECM, which could cause portions of the lumen to collapse. Given that Rab6 is part of the highly conserved “core” set of Rabs (Dunst et al., 2015), it is reasonable to assume that it is involved in maintaining the proper function of Golgi compartments also in TCs. Its depletion could thus cause a deficiency in the secretory system, and consequently an inability to secrete structural components of the aECM into the lumen. Rab2 has been associated with secretion in the fat body (Ke et al., 2018), consistent with an interpretation where this phenotype reflects a defunct secretory system. The defects that we found in *rab1* mutant TCs also included lumen gaps, consistent with a role of Rab1 in Golgi and secretory function (Charng et al., 2014; Sechi et al., 2017). The third feature, autofluorescent “fibres”, cannot be explained in this way. The deformation of the gas-filled portion of the lumen indicates that these fibres represent a solid or liquid material, and its autofluorescence suggests that it is the same material as the apical extracellular matrix itself. We speculate that this may be remnants of matrix from the previous larval moult that the TC failed to digest due to an inability to secrete the necessary enzymes.

The phenotypes that we observed in Rab8-depleted TCs are the most difficult to interpret. The hallmark of this phenotype was the formation of autofluorescent specks on the main tube, as well as smaller tubules that branch off from it and envelop it (“hairs”). This indicates a defect in the shaping of the apical membrane, as well as potentially an enlargement of the apical domain. Apical domain expansion can be induced by excess FGF signalling in TCs (Jones and Metzstein, 2011; Schottenfeld-Roames and Ghabrial, 2012), and the resulting phenotype is a curling up of the tube near the nucleus or near branch tips. The “postjunctional curls” and “curled tips” seen in Rab8-depleted TCs could thus be explained by this misregulation of domain size, but the “hairs” cannot. In other Drosophila epithelia, Rab8 plays a sorting role in polarised trafficking. Rab8-deficient cells secrete basal-directed cargo (Collagen) on their apical side (Devergne et al., 2017). Perhaps in TCs, basal structural or adhesive proteins missecreted on the apical side, i.e. in the subcellular tube, can cause invaginations from the tube that then also accumulate autofluorescent aECM (manifesting as specks) or extend and become tubules (“hairs”).

## Conclusion

We screened 26 Rab proteins to identify Rabs that are important for tracheal morphogenesis. Targeting a YFP-tag to knock down endogenously tagged Rab alleles, conferred high confidence that observed phenotypes are specific to the tagged Rab and not false positives. We identified 14 Rabs as required for dorsal branch morphogenesis during embryogenesis. The phenotypes indicated fate specification defects resulting in excess terminal cells, mismigration and failure of the fusion between contralateral dorsal branches. Some of the most severe phenotypes were associated with Rabs that are barely or not at all characterised, such as Rab19 and RabX5, making the tracheal dorsal branch a promising model to study their function. In contrast, only 4 Rabs were important for the morphogenesis of terminal cells during the larval stages. Rab2 and Rab6, were associated with abnormalities in the apical extracellular matrix that lines the lumen of the terminal cells’ subcellular tube network. These could be explained by defects in the secretory system, in which these Rabs are canonically involved. Rab8 and Rab10 were associated with terminal cell phenotypes that are hard to interpret because currently little is known about trafficking directed towards the basal membrane in terminal cells.

## Methods

### Drosophila strains and culture

All stocks were kept at 18°C and all crosses to generate lines were cultured at 25°C. All experimental crosses were cultured at 29°C to increase Gal4 expression. Only the crosses for YRab6 knockdown were cultured at 25°C because larvae did not reach the L3 stage at 29°C. The 27 YRab lines were a kind gift from M. Brankatschk (Dunst et al., 2015). *Btl-gal4* (Shiga et al., 1996) was recombined with a UAS-DsRed1 (BDSC 6282) construct on the third chromosome to create the driver line. To knock down the EYFP tag of the YRab alleles, we tested *GFP-IR1* (NIG-Fly, Mishima), *VALIUM20-EGFP.shRNA.3* (BDSC ID 41559) (Neumüller et al., 2012), and two deGradFP constructs (Caussinus et al., 2012), *P{UAS-Nslmb-vhhGFP4}2* (BDSC ID 38422) and *M{UASp-Nslmb.vhhGFP4}ZH-51D* (BDSC ID 58740). All four constructs are on the second chromosome. For the screen, we generated 27 lines carrying a YRab and GFP-IR1 and 27 lines carrying a YRab, *btl-gal4* and UAS-DsRed1. This required 10 recombined lines for Rabs on the second chromosome (Rab2, 3, 4, 5, 6, 9, 14, 30, 32, X1) and 10 recombined lines for Rabs on the third chromosome (Rab1, 7, 8, 11, 19, 23, 26, X4, X5, X6). We confirmed the presence of the YRab allele in all lines by PCR using genotyping primers flanking the start codon (see table S3), such that the product length increases if the YFP insertion just after the start codon is present. *Btl-gal4* was not homozygous viable and was balanced with *TM6b,Tb,Hu*. Likewise, GFP-IR1 was balanced with *CyO,dfdYFP* in some YRab-recombined lines. The Tb and dfdYFP markers were used to screen out balancer larvae during experiments.

For MARCM, we crossed males from an *hs-Flp; btl-gal4, UAS-GFP; FRT82B, tub-Gal80* driver line to virgins of the *FRT82B*, Rab1^S147213^/TM3 line (BDSC ID 37735) (Sechi et al., 2017). Embryos were collected for 6h at 25°C and then heatshocked at 38°C for 2h before being returned to 25°C culture. Third-instar wandering larvae were screened for GFP-positive terminal cells.

### Immunostaining for YRab expression

Third-instar wandering larvae of each YRab line were dissected according to standard protocol to expose the dorsal tracheal system and fixed for 15-20min in 8% paraformaldehyde. Fixed filets were kept in blocking buffer (1% w/v bovine serum albumin in phosphate-buffered saline) for 45min before incubation with anti-GFP antibody A-11122 (Thermo Fisher) diluted 1:500 in blocking buffer at 4°C overnight. Filets were then washed three times for 10min in PBS with 0.1% Triton-X and incubated with Alexa568-conjugated anti-rabbit secondary antibody diluted 1:500 in blocking buffer at room temperature for 1h. Filets were mounted in Vectashield medium with diamidino-phenylindole (DAPI; Vector Laboratories). We tested five different rabbit anti-GFP antibodies: A-11122 (Thermo Fisher), TP401 (Torrey Pines Biolabs), mAb anti-GFP (Upstate Biotec), Ab290 (Abcam) and antiGFP 598 (MBL). All five produced substantial unspecific staining in TCs of *w-*control larvae. This staining had a punctate distribution, similar to the distribution that Rabs typically show. We therefore quantified the staining intensity, comparing TCs of *yrab* larvae to *w*^*1118*^ larvae that were treated in parallel with the same batch of antibody.

### Heat-fixation for phenotype assessment

Third-instar wandering larvae of the respective cross were collected in distilled water with a brush, cleaned gently and then transferred to a coverslip with halocarbon oil 27 (Sigma). This was placed on a pre-heated heatblock at 65°C for 45s.

### Confocal imaging

All imaging was done on a Zeiss LSM780 inverted confocal laser scanning microscope equipped with a diode laser for 405nm excitation of tracheal extracellular matrix autofluorescence and DAPI, an Argon laser for 488nm excitation of GFP and YFP, and a DPSS 610-1 laser for 561nm excitation of DsRed, as well as a transmission photomultiplier tube detector to detect transmitted light. For terminal cell imaging, the objective used was a Plan-Apochromat 63X/1.4 Oil DIC M27 (Zeiss). For scoring dorsal tracheal anatomy phenotypes, a 20X Air Objective (Zeiss) was used as this allows a greater imaging depth, necessary to trace the sometimes very deep dorsal anastomoses.

### Computational analysis and phenotype scoring

To quantify YRab protein expression, a simple segmentation method was implemented in FIJI (Schindelin et al., 2012): A Gaussian blur filter was applied to the staining channel (561nm fluorescence). The resulting image was thresholded to obtain a mask of the terminal cell. A second region outside the terminal cell was chosen to measure background detector noise. The mean intensity in the background region was then subtracted from the mean staining fluorescence intensity of the cell region for each cell to obtain a proxy of staining level. The resulting values for each YRab were compared to the negative control cells recorded in the same microscopy session from the same staining experiment.

We created custom spreadsheets to record phenotype occurrences for each larva (respectively terminal cell; raw data in tables S1 and S2). We used the Shared Control, Two Groups and Multi Two Groups plots of the Estimation Stats online tool (Ho et al., 2018) for comparative estimation of effect sizes and p-value calculation. We considered effects significant if their 95% confidence interval was entirely above or below 0. Heatmap plots of phenotype frequencies were generated using the Morpheus online tool (Gould, 2016). Our initial screening for TC abnormalities included six additional phenotypes that we then discarded because they were overall too infrequent (“u-turn”, “cyst”, “triple junction”, “degenerate second lumen”), redundant (“hairy junction”), or their frequency never differed from controls (“wavy lumen”; can be seen in Fig. 4A).

## Data and reagent availability

The raw data (counts of phenotypes per larva, respectively per terminal cell) can be found in tables S1 and S2. Table S3 contains sequences of the genotyping primers we used to confirm the presence of the endogenous YFP-tag in the final *Drosophila* lines used for knockdown crosses. All data are available on Figshare (DOI to be confirmed).

## Acknowledgments

We thank M. Brankatschk for gifting us the YRab collection and the Advanced Light Microscopy Facility at EMBL for their support with imaging infrastructure. **Financial support**: B.B. was supported by the EMBL International PhD Programme.

## Supplemental data

**Figure S1.**
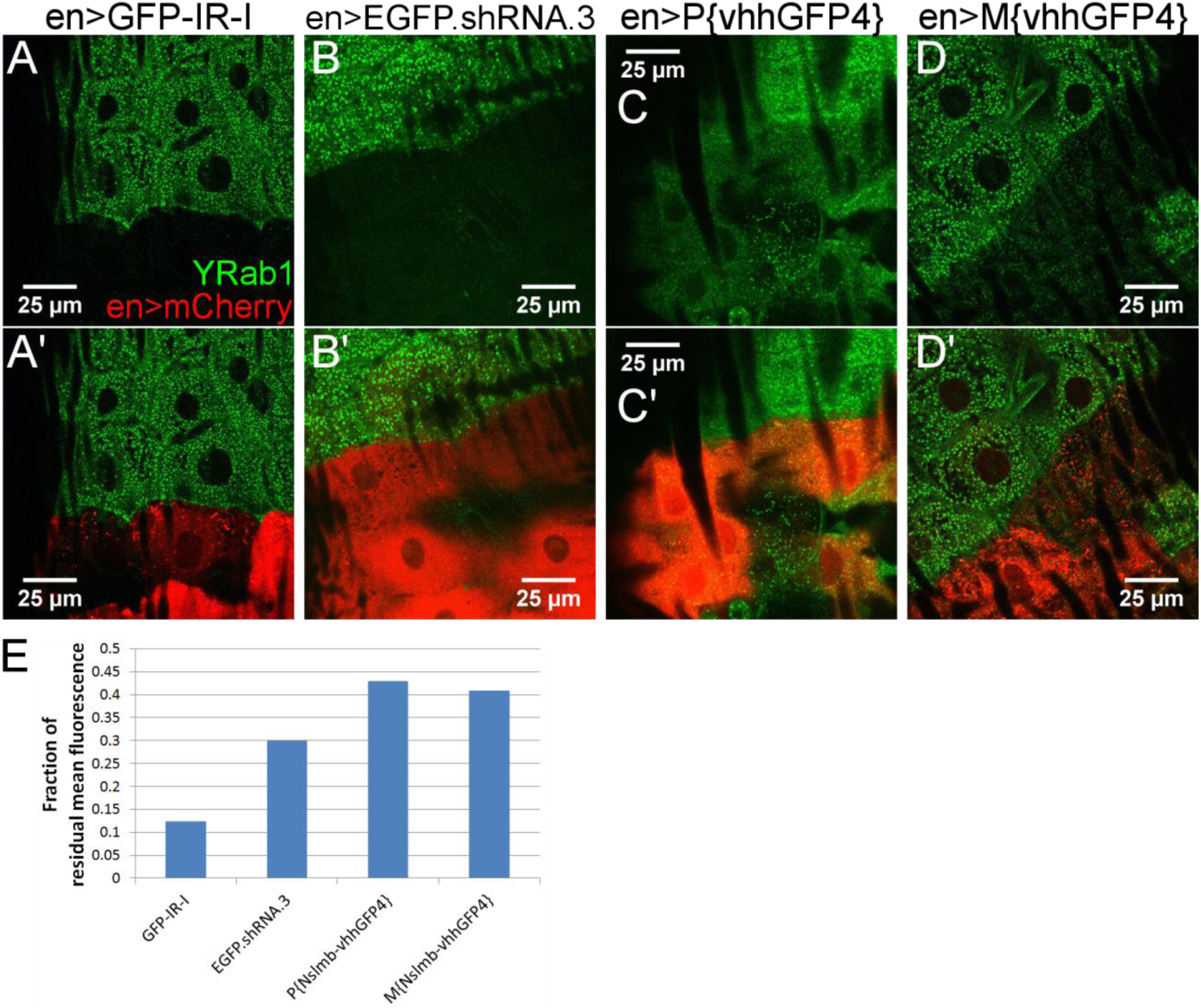
Comparison of YRab1 knockdown efficiency using four different knockdown constructs. We used the *engrailed*-gal4 driver to express each knockdown construct in a striped pattern in the epidermis of YRab1 larvae and imaged the endogenous YFP signal in heat-fixed third-instar wandering larvae. UAS-mCherry was used to label *engrailed*-gal4 expressing cells. (A-D’) Micrographs of epidermis in third-instar larvae showing a portion of epidermal cells not expressing *engrailed*-gal4 next to cells that express it (labelled by mCherry). (A-D) YFP signal from YRab1 (green), (A’-D’) Overlays of YRab1 with mCherry (red). (E) Quantification of knockdown efficiency. Bars show the difference in mean YRab1 pixel intensity in an area showing mCherry signal compared to YRab1 signal in an area showing no mCherry signal.

